# Comparative ‘omics analyses differentiate *Mycobacterium tuberculosis* and *Mycobacterium bovis* and reveal distinct macrophage responses to infection with the human and bovine tubercle bacilli

**DOI:** 10.1101/220624

**Authors:** Kerri M. Malone, Kévin Rue-Albrecht, David A. Magee, Kevin Conlon, Olga T. Schubert, Nicolas C. Nalpas, John A. Browne, Alicia Smyth, Eamonn Gormley, Ruedi Aebersold, David E. MacHugh, Stephen V. Gordon

## Abstract

Members of the *Mycobacterium tuberculosis* complex (MTBC) are the causative agents of tuberculosis in a range of mammals, including humans. A key feature of MTBC pathogens is their high degree of genetic identity, yet distinct host tropism. Notably, while *Mycobacterium bovis* is highly virulent and pathogenic for cattle, the human pathogen *M. tuberculosis* is attenuated in cattle. Previous research also suggests that host preference amongst MTBC members has a basis in host innate immune responses. To explore MTBC host tropism, we present in-depth profiling of the MTBC reference strains *M. bovis* AF2122/97 and *M. tuberculosis* H37Rv at both the global transcriptional and translational level via RNA-sequencing and SWATH mass spectrometry. Furthermore, a bovine alveolar macrophage infection time course model was employed to investigate the shared and divergent host transcriptomic response to infection with *M. tuberculosis* or *M. bovis*. Significant differential expression of virulence-associated pathways between the two bacilli was revealed, including the ESX-1 secretion system. A divergent transcriptional response was observed between *M. tuberculosis* and *M. bovis* infection of bovine alveolar macrophages, in particular cytosolic DNA-sensing pathways at 48 hours post-infection, and highlights a distinct engagement of *M. bovis* with the bovine innate immune system. The work presented here therefore provides a basis for the identification of host innate immune mechanisms subverted by virulent host-adapted mycobacteria to promote their survival during the early stages of infection.

**Importance:** The *Mycobacterium tuberculosis* complex (MTBC) includes the most important global pathogens for humans and animals, namely *Mycobacterium tuberculosis* and *Mycobacterium bovis,* respectively. These two exemplar mycobacterial pathogens share a high degree of genetic identity, but the molecular basis for their distinct host preference is unknown. In this work we integrated transcriptomic and proteomic analyses of the pathogens to elucidate global quantitative differences between them at the mRNA and protein level. We then integrated this data with transcriptome analysis of the bovine macrophage response to infection with either pathogen. Increased expression of the ESX-1 virulence system in *M. bovis* appeared a key driver of an increased cytosolic nucleic acid sensing and interferon response in bovine macrophages infected with *M. bovis* compared to *M. tuberculosis.* Our work demonstrates the specificity of host-pathogen interaction and how the subtle interplay between mycobacterial phenotype and host response may underpin host specificity amongst MTBC members.

## Introduction

The *Mycobacterium tuberculosis* complex (MTBC) comprises ten mycobacterial species that cause tuberculosis (TB) in a broad range of mammalian species, including humans (1-4). Typically, MTBC species show greater than 99% nucleotide sequence identity and yet exhibit distinct host preference, indicating that this low-level of genetic divergence holds major implications for host-pathogen interactions (1-3). Divergence in host tropism is illustrated through the comparison of the human adapted *Mycobacterium tuberculosis* with the animal bacillus *Mycobacterium bovis. M. tuberculosis* is a highly successful pathogen and is the world’s leading cause of death from an infectious agent with 1.7 million deaths reported in 2016 (5). *M. bovis* predominantly causes disease in cattle and bovine TB exacts a tremendous economic burden through production loss and control costs (6-8). *M. tuberculosis* appears unable to sustain (i.e., through cycles of infection, disease and transmission) in non-human animal populations, a fact that has been confirmed using an experimental bovine infection model (1, 9): while cattle infected with *M. bovis* display characteristic pathology, cattle infected with *M. tuberculosis* show minimal pathology despite positive skin-test and interferon-gamma responses indicative of successful infection. Conversely, while *M. bovis* can both infect humans and cause pulmonary disease that is clinically indistinguishable from *M. tuberculosis,* it rarely transmits among immunocompetent hosts (10, 11).

On a cellular level, the alveolar macrophage is the frontline host immune cell that encounters both *M. tuberculosis* and *M. bovis,* and its role during early stage infection is well established (12-18). Several studies have highlighted significant differences in the production of key innate factors, chemokines and cytokines at both the transcript and protein level in macrophages infected with *M. tuberculosis* or *M. bovis* (15-20). However, these studies evaluated only a subset of the innate response in macrophages and differences in the global transcript and protein response to infection with *M. tuberculosis* or *M. bovis* remains unknown. The central role of the alveolar macrophage during infection is also reflected in the fact that pathogenic mycobacteria have evolved several immune-evasion strategies to circumvent the killing mechanisms of the macrophage, including inhibition of phagosomal maturation, phagosomal escape and suppression of innate immune signalling (12-18). This facilitates the dissemination of the bacilli to other macrophages and ultimately throughout the host, with the concomitant development of immunopathology. Transmission of infection then occurs through the rupture of lesions into associated airways and the dispersal of bacilli (17, 18). Thus, it can be hypothesized that the initial interaction between host and pathogen may be key for the host preference observed between *M. tuberculosis* and *M. bovis;* whether this interaction has roots in host-centric or pathogen-centric processes, or indeed a combination of both, has yet to be fully elucidated.

*M. tuberculosis* H37Rv and *M. bovis* AF2122/97 were the first MTBC genomes to be fully sequenced and they represent the default reference strains for the human and animal tubercle bacilli (2, 21, 22). It is hypothesized that host tropism between these two species may be explained by differential gene expression profiles as a result of low genetic divergence (2, 21, 22). So far, functional studies have revealed that genetic changes between the two pathogens are responsible for differential nitrate reductase activity, for the loss of phenolic glycolipid production in *M. tuberculosis* H37Rv in contrast to *M. bovis,* and for differences in the PhoPR regulation system that governs the expression of virulence-related pathways such as EsxA/ESAT-6 secretion and cell wall lipid biosynthesis (23-26). While these studies highlight important differences between the two pathogens, host tropism likely involves a combination of events such as these that affect the expression and regulation of multiple virulence associated factors and/or the transcriptional regulators that govern their activity. In 2007, two microarray-based studies highlighted genes encoding the major antigens MPT83 and MPT70 that were expressed at higher levels in *M. bovis* and genes involved in SL-1 production that were expressed at a higher level in *M. tuberculosis* (27, 28). Since these reports, investigations into species-specific expression profiles of the two pathogens have been lacking at the global transcriptional level and have yet to be defined at the proteomic level. Definition of the differential “expressome” between *M. tuberculosis* and *M. bovis* will shed light on how alternate expression of two highly related genomes impacts on the ultimate success of these pathogens and host specificity within the MTBC.

As a route to defining host preference between *M. tuberculosis* and *M. bovis,* we have conducted in-depth profiling of *M. bovis* AF2122/97 and *M. tuberculosis* H37Rv at both the global transcriptional and translational level *in vitro* using RNA-seq and <u>S</u>equential <u>W</u>indow <u>A</u>cquisition of all <u>Th</u>eoretical Spectra (SWATH) mass spectrometry, a massively parallel targeting mass spectrometry that provides highly reproducible quantitative measurements across samples (29). To address how MTBC pathogen variation impacts on the host innate response, we have performed detailed comparative transcriptomic analyses of the bovine alveolar macrophage response to infection with both pathogens using RNA-sequencing (RNA-seq). Through these analyses, we reveal significant differential expression of virulence-associated pathways between *M. tuberculosis* and *M. bovis* was found, in particular the ESX-1 secretion system, while the macrophage infection study highlights a distinct engagement of *M. bovis* with the bovine innate immune system was found, in particular with the cytosolic DNA-sensing pathways of the macrophage.

## Methods

### Mycobacterial culture for pathogen transcriptomics and proteomics

Exponentially grown mycobacterial liquid cultures were established in Sauton’s basal media +0.025% tyloxapol. For mid to late-log phase culture, mycobacterial cells were grown to an optical density (OD_600nm_) of 0.6 - 0.8 at 37°C prior to harvest. For the current study, six *M. bovis* AF2122/97 and six *M. tuberculosis* H37Rv replicates were prepared. Matched RNA and protein samples were harvested and prepared for strand-specific RNA-sequencing or SWATH mass spectrometry.

### RNA extraction, RNA-seq library preparation and high-throughput sequencing for *M. bovis* and *M. tuberculosis*

Mycobacterial cells were harvested by centrifugation at 2,500 × g for 10 min and the pellet was re-suspended in 1 ml of TRIzol^®^ (Life Technologies). The suspension was transferred to a 2 ml screw cap tube and the cells were lysed by bead-beating for 30 s at maximum setting using 1 mm glass beads (Sigma) on a MagNaWLyser instrument (Roche). Samples were placed at 80°C immediately and thawed before use. 20% v/v chloroform was added, the sample were shaken vigorously for 15 s and incubated for 2-3 min at room temperature. The samples were centrifuged at 12,000 × g for 15 min at 4°C and the top phase was added to the DNA-free columns from the RNeasy plus kit (Qiagen). The sample were processed as per the manufacturer’s guidelines with the following exceptions: 1.5 volumes of 100% ethanol was added to sample prior to its application to the RNeasy column in order to recover all RNA species. RNA was eluted in molecular grade water and its concentration was determined using the NanoDrop spectrophotometer (NDW1000) prior to DNase treatment. A DNase treatment using TURBO DNase kit (Thermo Fisher Scientific) was performed by following the vigorous DNase treatment as per manufacturer’s guidelines; 7 μl of DNase buffer + 1 μl enzyme at 37°C for 30 min followed by a further 1 μl of enzyme and incubation at 37°C for 30 min. A DNase stopping solution was not added to the samples as they were column-purified and concentrated using the RNA Clean and Concentrator kit according to manufacturer’s guidelines (Zymo). RNA was eluted in molecular grade water and its concentration was determined using the NanoDrop spectrophotometer (NDW1000). RNA integrity number (RIN) values were assessed for each RNA sample being considered for RNA-sequencing using the 2100 Bioanalyser (Agilent) and the RNA 6000 Nano kit (Agilent) according to manufacturer’s guidelines. RIN values are calculated by assessing the entire electrophoretic trace of an RNA sample, along with the 23S/16S rRNA intensity value. Only samples with a RIN value > 8 were selected for further analysis by RNA-sequencing. Sequencing libraries were prepared at the Genomics Core, Michigan State University, Michigan, USA using the Illumina Truseq Stranded Total RNA Library Prep Kit LT and the Epicenter RiboWZero Magnetic Bacteria kit to deplete ribosomal RNA. Single-end, strand-specific 50 bp read data was produced with base calling performed by Illumina Real Time Analysis (RTA) v1.18.64.

### Differential gene expression analysis of *M. bovis* and *M. tuberculosis* RNA-sequencing data

Computational analyses were performed on a 32-node Compute Server with Linux Ubuntu [version 12.04.2]. Briefly, adapter sequence contamination and paired-end reads of poor quality were removed from the raw data. At each step, read quality was assessed with FastQC [version 0.10.1] (30). Single-end reads were aligned to the *M. bovis* AF2122/97 or *M. tuberculosis* H37Rv reference genomes with the aligner Stampy in hybrid mode with BWA (31). Read counts for each gene were calculated using featureCounts, set to unambiguously assign uniquely aligned single-end reads in a stranded manner to gene exon annotation (32).

Prior to cross-species differential expression analysis, an Identical/Variable gene dataset was constructed for *M. tuberculosis* H37Rv and *M. bovis* AF2122/97 where orthologous genes were separated into those genes whose protein products are of equal length and 100% conserved at the amino acid level (Identical, *n =* 2,775) from those that are not (Variable, *n =* 1,224). (Fig.1A, Supp_I.xlsx). Among the Variable genes are examples of truncated genes, genes that have been split into two or more as a result of in-frame sequence variance (leading to some genes being represented in more than one Variable gene pair), or genes that differ by a non-synonymous SNP resulting in an amino acid change at the protein level. Negative binomial modelling tools such as DESeq2 that was used in this instance assume equal feature lengths when calculating differential expression (DE) of a gene, or in this case between orthologous genes of two species in a given condition (33). For those annotations whose gene lengths are not equal, such as in the case of truncated/elongated/frameshift instances found in the *M. bovis* AF2122/97 genome with respect to *M. tuberculosis* H37Rv, analysis with DESeq2 would result in erroneous differential expression results; thus a separate differential expression analyses was carried out for Variable genes using <u>T</u>ranscript per <u>M</u>illion (TPM) values that are normalised for feature length (33, 34). Differential gene expression analysis for those genes of equal lengths was performed using the DESeq2 pipeline, correcting for multiple testing using the Benjamini-Hochberg method (33). All further reference to differentially expressed genes between the two mycobacterial species will be with regards to a gene being expressed at a higher level in one species with respect to the other, and hence if a gene is upregulated in *M. bovis* it is downregulated or expressed at a lower level in *M. tuberculosis* and vice versa (log_2_fold change (Log_2_FC > 1 and < -1), false discovery rate (FDR) threshold of significance < 0.05).

**Figure 1:**
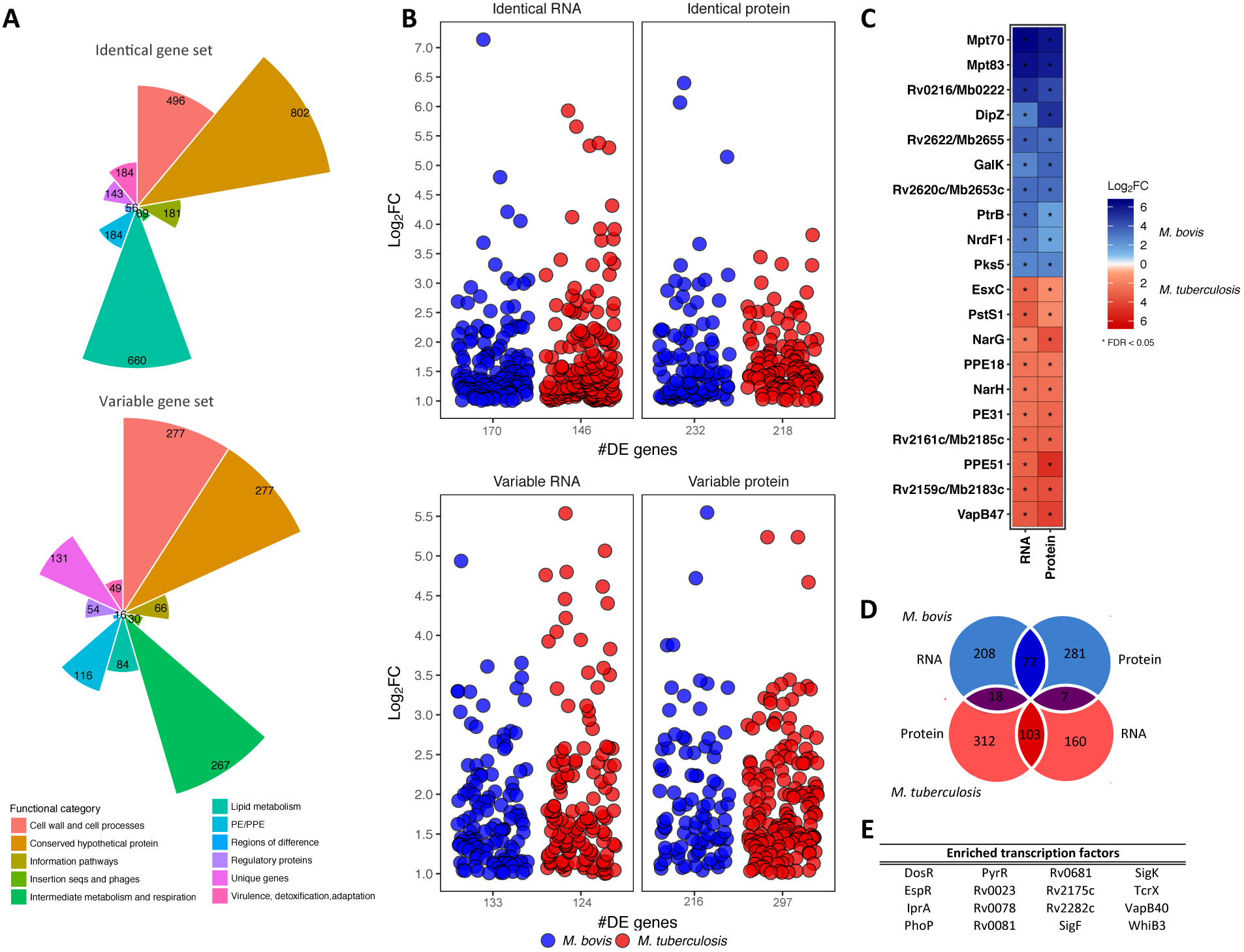
**A)** The number of genes from *M. bovis* AF2122/97 and *M. tuberculosis* H37Rv classified as either “Identical” (100% conserved in length and amino acid sequence, top plot) or Variable (all other orthologous genes, bottom plot). Colours represent the various gene categories to which each gene belongs. B) The level of expression (‘Log_2_FC’) of Identical genes (top panel) and Variable genes (bottom panel) that are differentially expressed (|Log_2_FC| > 1, FDR < 0.05) and upregulated in either *M. bovis* (blue) or *M. tuberculosis* (red) at both the RNA and protein level. The number of genes in each category is indicated on the x-axis (‘æDE genes’). C) The top 20 differentially expressed genes at the RNA and protein level (|Log_2_FC| > 1, FDR < 0.05 (‘*’)) that are upregulated in *M. bovis* (blue) or *M. tuberculosis* (red). D) The overlap of genes that are upregulated in either *M. bovis* (blue) or *M. tuberculosis* (red) at the RNA and protein level. Dark blue overlap represents those genes upregulated in *M. bovis* only, dark red overlap represents those genes upregulated in *M. tuberculosis* while purple overlaps represent those genes that show discordant expression patterns at the RNA and protein level between the two species. **E)** The transcription factors enriched for Identical genes (100% conserved in length and amino acid sequence between the two species) that are differentially expressed between *M. bovis* and *M. tuberculosis.* (Rand Index, *P <* 0.01).

### Transcription factor enrichment analysis

Data relating to the shift in the transcriptional landscape of *M. tuberculosis* upon overexpression of 183 transcription factors was used to perform a formal transcription factor enrichment analysis (35-37). The data represents 9,335 regulatory events and provides regulatory evidence for over 70% of the annotated genes in the *M. tuberculosis* genome (FC > 2, *P <* 0.01) (35-37). This data was analysed alongside the DE genes identified in this study between *M. bovis* and *M. tuberculosis.* Only genes and transcription factors that are 100% identical in sequence and length between the two species were considered for this analysis. Over-representation of a transcription factor with a given set of differentially expressed genes was assessed by gene-regulon association and calculation of the Rand Index (Log_2_FC > 1 and *P <* 0.05 for a given DE gene).

### Protein extraction and SWATH mass spectrometry for *M. bovis* and *M. tuberculosis*

Mycobacterial cells were harvested by centrifugation at 2,500 × *g* for 10 min and the pellet was re-suspended in 1 ml LB buffer (0.1 M ammonium bicarbonate buffer, 8 M urea, 0.1% *Rapi*GEST SF (Waters, UK)). The suspension was transferred to a 2-ml screw cap tube and the cells were lysed by bead-beating for 30 s at maximum setting using 3 mm glass beads (Sigma) and a MagNaLyser instrument (Roche). The lysate supernatant was harvested by centrifugation at 12,000 x *g* for 10 min and transferred to a clean 1 ml tube. The remaining pellet was re-suspended in LB buffer and the bead beating cycle was repeated twice more. Protein lysate samples were stored at -80°C. Protein samples were removed from -80°C storage and thawed on ice. Total protein content was measured using the Qubit Protein Assay kit according to manufacturer’s guidelines and protein concentrations were adjusted to 0.5 mg/ml. Protein disulphide bonds were reduced by addition of 0.2 M Tris(2-carboxyethyl)phosphine (TCEP) and the resulting free cysteine residues were alkylated by addition of 0.4 M iodoacetamide (IAA). Extracted protein samples were diluted with 0.1 M ammonium bicarbonate buffer to reach a urea concentration of < 2 M and then digested with 1:50 enzyme/substrate ratio of sequencing grade modified trypsin (Promega). 50% trifluoroaceticacid (TFA) was added to lower the pH to 2 in order to stop the tryptic digest and to precipitate the *Rapi*GEST. Water-immiscible degradation products of *Rapi*GEST were pelleted by centrifugation at 12,000 x *g* for 10 min. The cleared peptide solution was desalted with C18 reversed-phase columns (SepWPak Vac C18, Waters). The columns were pre-conditioned 2-3 times with acetonitrile and equilibrated 3 times with Buffer A (2% acetonitrile, 0.1% trifluoroacetic acid in H_2_O) prior to sample loading. The flow-through was re-loaded onto the column and the column was then washed 3 times with Buffer A. The peptides were eluted from the column using Buffer B (50% acetonitrile, 0.1% trifluoroacetic acid in H_2_O) and the elution step was repeated. The eluate was dried under vacuum using a rotary evaporator at 45°C. Dried peptide pellets were re-suspended in MS buffer (2% acetonitrile, 0.1% trifluoroacetic acid in ultra pure H_2_O) to a concentration of 1 μg/μl, sonicated in a water bath for 3 min and supernatant was harvested by centrifugation at 12,000 *×g* for 10 min.

SWATH mass spectrometry measurements were conducted at the Institute for Molecular Systems Biology at ETH Zurich. 1 μg of each peptide sample was measured in SWATH mode on a TripleTOF 5600 mass spectrometer using data-independent acquisition settings as described earlier (29, 38-40). Resulting raw SWATH data was analysed using an automated pipeline and the software OpenSWATH with the *M. tuberculosis* H37Rv SWATH assay library (38). Differential expression analysis of protein identified in *M. tuberculosis* and *M. bovis* samples was dependent on the detection of the protein in both species. The difference in protein fold changes and the corresponding FDR corrections between *M. tuberculosis* and *M. bovis* were calculated using MSstats (39, 41). A |Log_2_FC| > 0.56 and an FDR < 0.05 was required for a protein to defined as differentially expressed between *M. tuberculosis* and *M. bovis.*

### Animals

All animal procedures were performed according to the provisions of the Irish Cruelty to Animals Act of 1876 with ethical approval from the University College Dublin (UCD) Animal Ethics Committee (protocol number AREC-13-14-Gordon). Ten unrelated Holstein-Friesian male calves (7-12 weeks old) were maintained under uniform housing conditions and nutritional regimens at the UCD Lyons Research Farm (Newcastle, County Kildare, Ireland). All animals were selected from a tuberculosis-free herd that is screened annually using the single intradermal comparative tuberculin skin test.

### Alveolar macrophage isolation, cell culture and infection

The laboratory methods used to: (1) isolate, culture and infect bovine alveolar macrophages with *M. bovis* and *M. tuberculosis,* and (2) generate strand-specific RNA-seq libraries using RNA harvested from these cells has been described in detail by us elsewhere (15, 42). An abridged description of the laboratory methods used in this study is provided below and the complete bioinformatics pipeline is accessible online (https://github.com/kerrimalone/AlvMac). Total lung cells were harvested by pulmonary lavage of lungs obtained post-mortem and stored in freezing solution (10% DMSO (Sigma-Aldrich Ltd.), 90% FBS) at a density of 2.5 × 10^7^ cells/ml in 1 ml cell aliquots at -140°C. When required, the cell pellet was resuspended in 15 ml of R10^+^ media and placed in a 75 cm^2^ vented culture flask (CELLSTAR®, Greiner Bio-One Ltd.) and incubated for 24 h at 37°C, 5 % CO_2_. After incubation, media was removed together with non-adherent cells, adherent cells were washed with 15 ml HBSS pre-warmed to 37°C and dissociated by adding 10 ml pre-warmed 1× non-enzymatic cell dissociation solution (Sigma-Aldrich Ltd.) to each culture flask. Cells were then pelleted (200 × g for 5 min), resuspended in 10 ml pre-warmed R10+ media and counted using a Vi-CELL™ XR Cell Viability Analyzer and reagent kit (Beckman Coulter Inc.). Mean viable cell recovery was estimated at ∼ 80% for each animal. Cell counts for each animal were adjusted to 5 × 10^5^ cells/ml (based on viable cell counts) using pre-warmed R10+ media, seeded at 5 × 10^5^ cells/well on individual 24-flat well tissue culture plates (Sarstedt Ltd.) and incubated for a further 24 h at 37°C, 5 % CO_2_, until required for mycobacterial infection. The purity of the seeded macrophages for each animal samples was 95% as estimated by flow cytometry analysis (data not shown).

*M. bovis* AF2122/97 and *M. tuberculosis* H37Rv were cultured in Middlebrook 7H9-ADC medium containing either 0.2% v/v glycerol for *M. tuberculosis* or 10 mM sodium pyruvate for *M. bovis* at 37°C until mid-logarithmic phase. Prior to infection, mycobacterial cultures were pelleted by centrifugation (200 × g, 10 min), pellets were disrupted with 3 mm sterile glass beads (Sigma-Aldrich Ltd.) by vortexing at top speed, 1 min. Cells were resuspended in pre-warmed R10 media, sonicated at full power (Branson Ultrasonics Corporation) for 1 min and the cell number was then adjusted to 5 × 10^6^ bacterial cells/ml (OD_600nm_ of 0.1 = 1 × 10^7^ bacterial cells) for a multiplicity of infection (MOI) of 10 bacilli per alveolar macrophage.

For the infection time course, the R10 media was removed from the macrophages and replaced with 1 ml R10 media containing *M. bovis* or *M. tuberculosis* (5 × 10^6^ bacilli/ml); parallel non-infected control alveolar macrophages received 1 ml R10 media only. The alveolar macrophages were incubated at 37°C, 5 % CO_2_ for times of 2, 6, 24 and 48 hours post-infection (hpi). Following completion of the 2 hpi time point, the 2 hpi macrophages were lysed (by adding 250 μl RLT-1% β-mercaptoethanol buffer per tissue culture plate well) and stored at -80°C, while the media for the 6, 24 and 48 hpi macrophages was replaced with 1 ml fresh RIO media per well and cells were reincubated at 37°C, 5 % CO_2_ until required for harvesting. CFU were monitored over the infection time course (Fig.S1).

### RNA extraction, RNA-seq library preparation and high-throughput sequencing for bovine alveolar macrophage samples

For the current study, 117 strand-specific RNA-seq libraries were prepared. These comprised *M. bovis-, M. tuberculosis-* and non-infected samples from each time point (0, 2, 6, 24 and 48 h) across 10 animals (with the exception of one animal that did not yield sufficient alveolar macrophages for *in vitro* infection at the 48 hpi time point). RNA extractions from macrophage lysates included an on-column genomic DNA elimination step (RNeasy^8^ Plus Mini kit (Qiagen Ltd)). RNA quantity and quality was assessed using a NanoDrop™ 1000 spectrophotometer (Thermo Fisher Scientific Inc.) and a Bioanalyzer and an RNA 6000 Nano LabChip kit (Agilent Technologies Ltd). All samples displayed 260/280 ratio > 2.0 and RNA integrity numbers > 8.5. 200 ng total RNA from each sample was used for RNA-seq library preparation. Poly(A) mRNA enrichment was performed (Dynabeads^®^ mRNA DIRECT™ Purification Kit (Invitrogen, Life Technologies)) and Poly(A)-enriched mRNA was used to prepared individually barcoded strand-specific RNA-seq libraries (ScriptSeq™ version 2 RNA-Seq Library Preparation Kit (Illumina, San Diego, CA, USA)). The libraries were pooled into three sequencing pools and sequenced across 24 flow cell lanes (lllumina^®^ HiSeq2000, Beijing Genomics Institute, Shenzhen, China).

### Differential gene expression analysis of bovine alveolar macrophage RNA-sequencing data

Computational analyses was performed on a 32-node Compute Server with Linux Ubuntu [version 12.04.2]. Briefly, pooled libraries were deconvoluted, adapter sequence contamination and paired-end reads of poor quality were removed. At each step, read quality was assessed with FastQC [version 0.10.1] (30). Paired-end reads were aligned to the *Bos taurus* reference genome (*B. taurus* UMD3.1.1) with STAR aligner (43). Read counts for each gene were calculated using featureCounts, set to unambiguously assign uniquely aligned paired-end reads in a stranded manner to gene exon annotation (32). Differential gene expression analysis was performed using the edgeR Bioconductor package that was customized to filter out all bovine rRNA genes, genes displaying expression levels below one count per million [CPM] in at least ten individual libraries and identify differentially expressed (DE) genes between all pairs of infection groups within each time point, correcting for multiple testing using the Benjamini-Hochberg method with Log_2_FC > 1 and < -1 and an FDR threshold of significance < 0.05 (44, 45). Cellular functions and pathways over-represented in DE gene lists were assessed using the SIGORA R package (46).

## Data availability

RNA-seq datasets can be found using accession number PRJEB23469. SWATH MS data and OpenSWATH outputs can be found on PeptideAtlas under identifier PASS00685 (http://www.peptideatlas.org/PASS/PASS00685).

## Results

### Differential RNA and protein expression between *M. bovis* and *M. tuberculosis*

For this study, 12 strand-specific RNA-seq libraries were prepared from *M. bovis* AF2122/97 (*n =* 6) and *M. tuberculosis* H37Rv (*n =* 6) grown exponentially in Sauton’s basal media pH 7.0 (Fig.S2; mapping statistics can be found in Supp_II.csv). An ‘Identical’/’Variable’ gene dataset was constructed where orthologous genes between the two species were separated into those genes whose protein products are of equal length and 100% conserved at the amino acid level (Identical, *n =* 2,775) from those that are not (Variable, *n =* 1,224). (Fig.1A, Supp_I.xlsx). 170 and 146 differentially expressed (DE) Identical genes and 133 and 124 DE Variable genes were identified for *M. bovis* and *M. tuberculosis* respectively, amounting to 573 DE genes in total (Fig.1A, B, Supp_III.xlsx). Twelve SWATH mass spectrometry (MS) datasets were generated from total protein samples harvested from the same cultures as the RNA (Fig.S2, Fig.S4, Supp_III.xlsx). Overall, 2,627 proteins were detected using the *M. tuberculosis* assay library (∼70% and ∼56% of total Identical and Variable protein, respectively) (Fig.S4, Supp_III.xlsx) (38, 40). Of the 1,937 Identical proteins detected by SWATH MS, 232 and 218 proteins were found to be upregulated *M. bovis* and *M. tuberculosis* respectively, totalling 450 DE proteins (Fig.1A, B, Supp_III.xlsx). 133 and 215 Variable proteins were found to be upregulated *M. bovis* and *M. tuberculosis* respectively, amounting to 348 DE Variable proteins in total (50.4%) (Fig.1A, B, Supp_III.xlsx). Overlap of the DE lists for *M. bovis* and *M. tuberculosis* revealed 77 and 103 genes that are significantly upregulated in either species at both the RNA and protein level respectively (Fig.1D); the top 20 of these are represented in Fig.1C.

Genes encoding Mpt70 and Mpt83 are the top two genes upregulated at the RNA and protein level in *M. bovis;* this is a result of a non-operational anti-SigK protein in *M. bovis* leading to constitutive upregulation of the SigK regulon, of which the *mpt83* and *mpt70* genes are components (47). Furthermore, Rv0216/Mb0222, a double hotdog hydratase, is upregulated in *M. bovis* at both the RNA and protein level as previously observed by microarray analysis (27). Amongst the genes upregulated at both the RNA and protein level in *M. tuberculosis* H37Rv are: *ppe51;* antitoxin *vapB47;* and nitrate reductase associated genes *narH* and *narG,* previously reported as upregulated at the RNA level in *M. tuberculosis* in comparison to *M. bovis* as a result of a SNP in the promoter region of *narGHJI* (Fig.1C)(25, 26). Incomplete overlap between DE genes at the transcriptional and translational level seen in this study has been reported in other studies and can be attributed to post-transcriptional and post-translational regulation within the cell, but also to more technical aspects, such as differences in detection limits and particular thresholds chosen to define differentially expressed RNA or protein (48-50).

### Transcription factor enrichment analysis: the PhoP regulon and ESX-1 secretion system

The differential gene expression observed between *M. bovis* and *M. tuberculosis* may be a consequence of differences in global transcriptional network regulation between the two species. To address this hypothesis, a formal transcription factor enrichment analysis was performed and revealed the significant association of 16 transcription factors with the DE Identical genes between *M. bovis n* = 146) and *M. tuberculosis* (*n* = 170) (Fig.1E) (37). The association of transcription factors such as alternate sigma factors SigK and SigF along with cytoplasmic redox sensor WhiB3 with the DE gene lists indicates that disparate expression of virulence-related pathways regulated by these transcription factors between the two pathogens could have significant consequence for infection (47, 51, 52). Furthermore, PhoP, EspR and DosR are also significantly associated with the DE genes; these transcription factors are important for adaptation of *M. tuberculosis* to the intracellular environment and are functionally linked by such processes (Fig.1E) (53, 54).

The PhoPR two-component system has a major role in regulating the pathogenic phenotype of *M. tuberculosis* by controlling the expression of a variety of virulence-associated pathways including SL-1, DAT and PAT lipid production and the Type-VII secretion system ESX-1; mutations in the PhoPR system of *M. bovis* have been suggested to play a role in the host specificity between the bovine- and human-adapted mycobacterial species (23, 53, 55-57). Further investigation into the 72-gene regulon of PhoP identified 33 DE genes between the two species and these are presented in Fig.2A. The production of lipids SL-1 and PDIM is under PhoP regulation and is coupled within the *M. tuberculosis* cell; intriguingly *M. bovis* is reported to lack SL-1 in the cell envelope (58-60). In this study, the expression of genes associated with the biosynthesis of SL-1 (*e.g. papA1, papA2, pks2, mmpL8*) was at a higher level in *M. tuberculosis* and conversely, genes associated with the biosynthesis of PDIM (*e.g. ppsA-E, lppX*) were expressed at a higher level in *M. bovis* at the RNA and protein level (Fig.S5). SL-1 is one of the most abundant lipids in the mycobacterial outer membrane, is unique to pathogenic mycobacteria, is immunogenic, and is implicated in the alteration of phago-lysosome fusion. Likewise, PDIM is required for mycobacterial virulence, facilitates macrophage invasion and protects against reactive nitrogen species. The differential expression of lipid-associated systems between the two species therefore presents distinct lipid-repertoires to interact with the host that could affect the overall success of infection (61-67).

**Figure 2:**
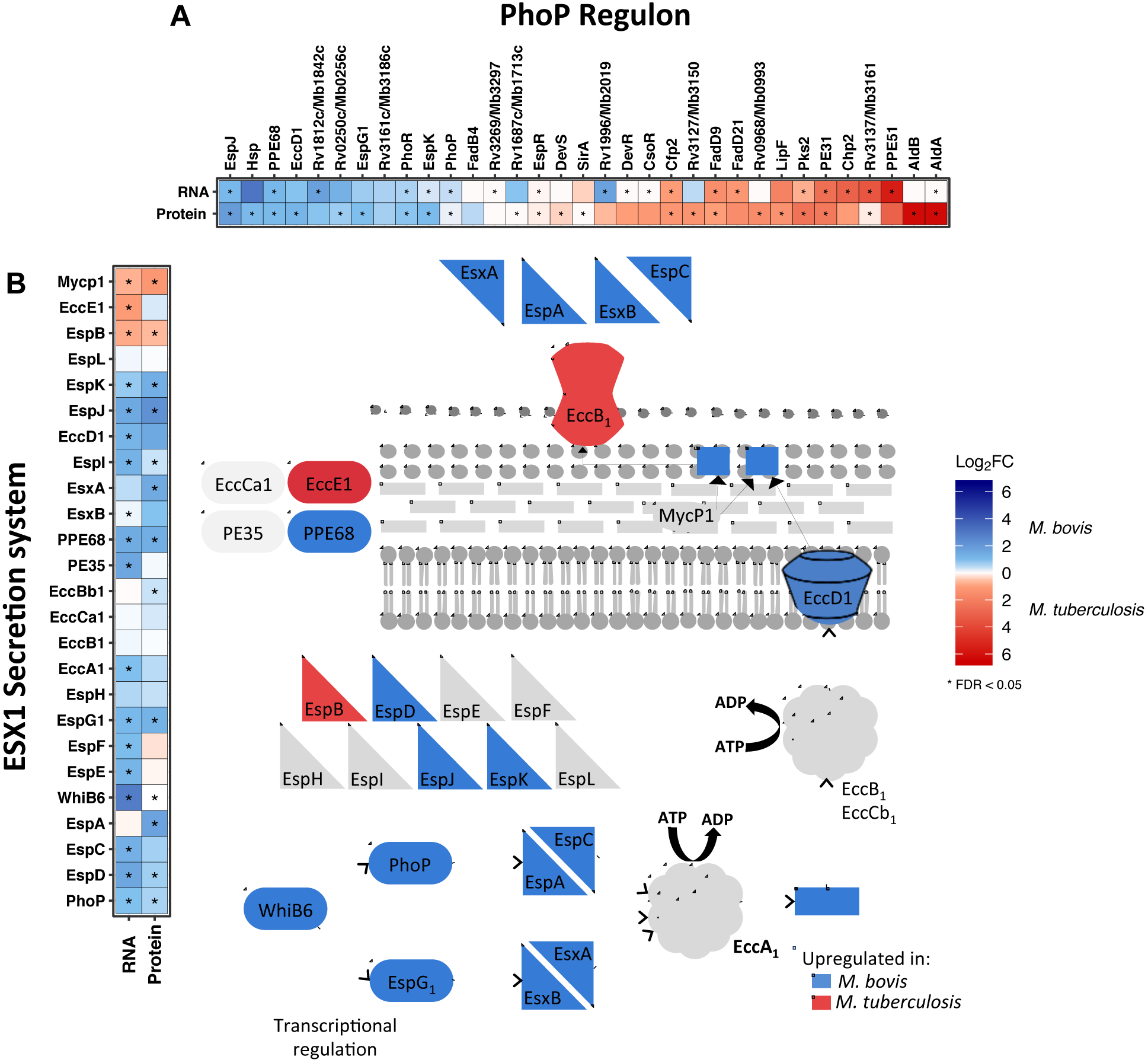
The differentially expressed genes (|Log_2_FC| > 1, FDR < 0.05 (‘*’)) belonging to A) the PhoP regulon and B) the ESX-1 secretion system that are upregulated in *M. bovis* (blue) or *M. tuberculosis* (red). Inset is a representation of the ESX-1 secretion system pathway of *M. tuberculosis* coloured according to the upregulation of the associated gene in *M. bovis* (blue) or *M. tuberculosis* (red).

The major antigens ESAT-6 and CFP10 are secreted by the ESX-1 secretion system of *M. tuberculosis,* a system which has been implicated in mycobacterial escape from the phagosome to the cytosol that results in a Type-I interferon response within the infected macrophage (57, 68, 69). PhoP and EspR regulate the expression of ESX-1 secretion system-related genes and as stated are significantly associated with the DE genes between the two pathogens; despite EspR being expressed to a higher level in *M. tuberculosis* (Supp_III.xlsx), there is a significant upregulation of the ESX-1 secretion system in *M. bovis* in comparison to *M. tuberculosis,* including ESX-1-related proteins such as EsxA, EspA, EspC, EspD at both the transcriptional and translational level (55-57, 70, 71) (Fig.2B). Furthermore, PhoP was expressed to a higher level in *M. bovis;* this may represent an attempt at a compensatory mechanism for aberrant PhoP signalling and supports previous reports of suboptimal PhoP signalling in *M. bovis* (23, 59). Seven of 55 genes regulated by DosR were expressed higher at the RNA level in *M. tuberculosis,* likely reflecting the intimate relationship of DosR and its associated regulon with that of PhoP/EspR/WhiB3 (Supp_IV.xIsx).

### A “core” macrophage response common to infection with either species

117 strand-specific RNA-seq libraries were prepared from bovine alveolar infected macrophages that comprised *M. bovis-, M. tuberculosis-* and non-infected samples from each time point (0, 2, 6, 24 and 48 hpi) across 10 animals, with the exception of one animal that did not yield sufficient alveolar macrophages for *in vitro* infection at the 48 hour post-infection time point). Matched non-infected macrophage control samples were included for all infection time-points (Fig.S2). Quality control and mapping statistics can be found in Supp_V.csv and Fig.S5.

The comparison of *M. bovis-* or *M. tuberculosis-infected* macrophages with respect to non-infected macrophages revealed a sequential increase in the number of DE genes across the infection time-course, which peaked at 48 hpi and a larger number of DE genes were seen in *M. bovis*-infected macrophages with the exception of 6 hpi (Fig.3A, B, C, Supp_VI.csv); similar temporal expression profiles were previously reported in other *in vitro* bovine and human macrophage infection studies (42, 72-74). Comparison of these DE gene lists identified a subset of genes that displayed the same directionality and a similar magnitude of expression (Fig.4A) (Supp_VII.csv). The association of enriched pathways such as *Cytokine-cytokine receptor interaction, NOD-like receptor signalling* and *Jak-STAT signalling* with this gene subset suggests a robust “core” macrophage response to infection with either mycobacterial species throughout the time course (Fig.4B). The core response includes numerous key genes known to be involved in the innate immune response against pathogenic mycobacteria such as: *CCL20* (75); *IL18,* which limits the growth of *M. tuberculosis* in human macrophages (76-78); anti-inflammatory *IL10* (79); and *NOS2,* polymorphisms of which are associated with susceptibility of Holstein cattle to bovine TB (80) (Fig.4C). Furthermore, the *HIF-1* signalling pathway was significantly enriched for the DE genes common to both infection series; this pathway is associated with regulating a switch in central glucose metabolism during high-energy demanding events, such as infection, in neutrophils and macrophages (81)(Fig.S6A).

**Figure 3:**
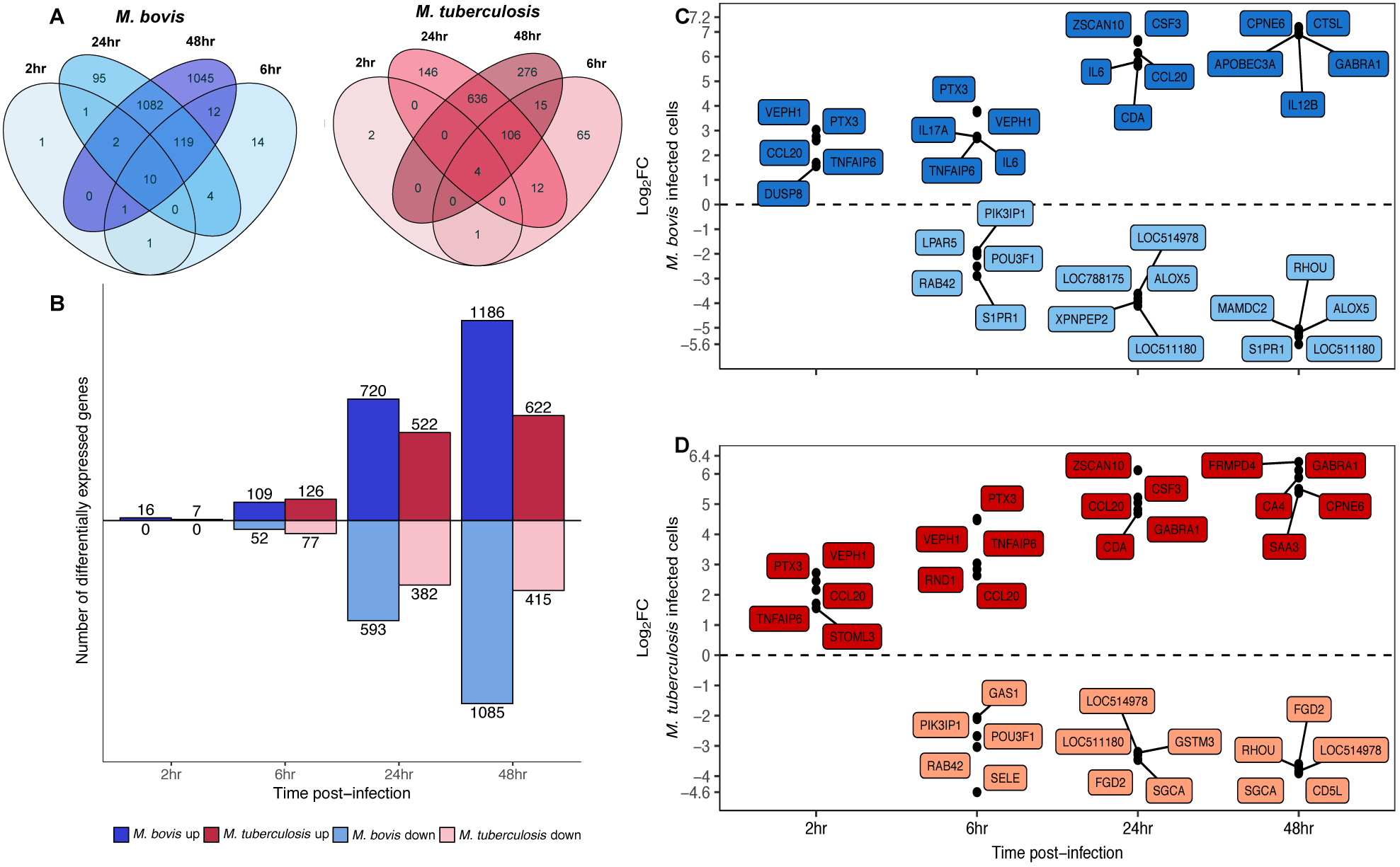
**A)** The differentially expressed genes (|Log_2_FC| > 1, FDR < 0.05) of bovine alveolar macrophages infected with *M. bovis* (blue) or *M. tuberculosis* (red) at 2, 6, 24 and 48 hours post-infection. **B)** The number (y-axis) and direction of change (up = positive y-space, down = negative y-space) of differentially expressed genes (|Log_2_FC| > 1, FDR < 0.05) of bovine alveolar macrophages infected with *M. bovis* (blue) or *M. tuberculosis* (red) at 2, 6, 24 and 48 hours post-infection (x-axis). **C)** The top 5 upregulated (positive y-space) and 5 downregulated (negative y-space) differentially expressed genes (|Log_2_FC| > 1, FDR < 0.05) of bovine alveolar macrophages infected with *M. bovis* or **D)** *M. tuberculosis* at *2,* 6, 24 and 48 hours post-infection (x-axis).

**Figure 4:**
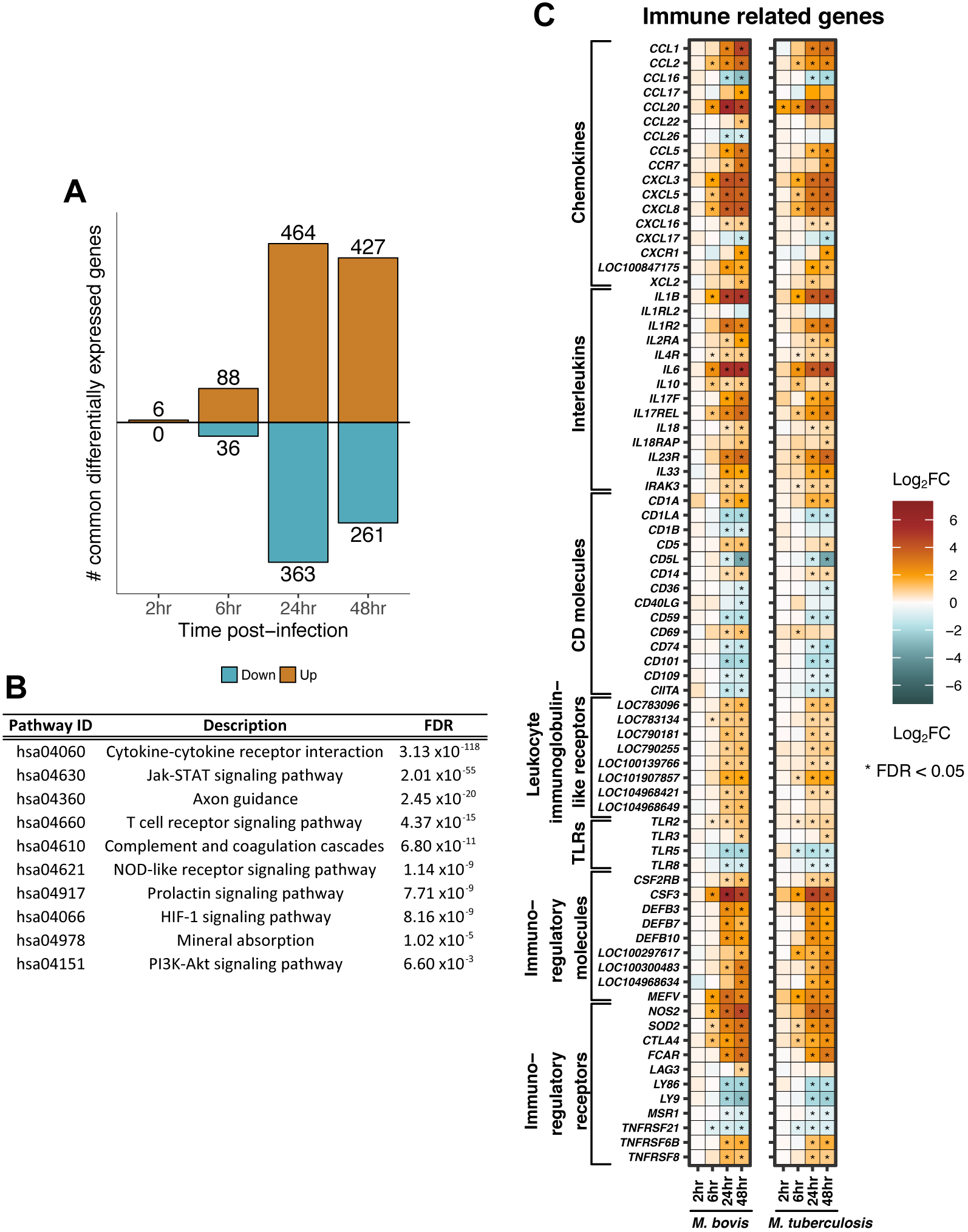
**A)** The number (y-axis) and direction of change (up = orange, down = cyan) of genes that are commonly differentially expressed (“core response”) (| Log_2_FC | > 1, FDR < 0.05, with non-significant delta comparison values) in bovine alveolar macrophages infected with *M. bovis* and infected with *M. tuberculosis* at 2, 6, 24 and 48 hours post-infection. B) Pathways enriched for 688 genes that are commonly differentially expressed (“core response”) in bovine alveolar macrophages infected with *M. bovis* and *M. tuberculosis* over the first 24 hours of infection (FDR < 0.05). **C)** Genes that are commonly differentially expressed (“core response”) (|Log_2_FC| > 1, FDR < 0.05 (‘*’)) and associated with the innate immune response in bovine alveolar macrophages infected with *M. bovis* (left column) or *M. tuberculosis* (right column) over 48 hours post-infection.

### DNA sensing and RIG-I like signalling pathways are found in the divergent response to infection with *M. tuberculosis* and *M. bovis*

Aside from the defined “core” response genes, there were larger numbers of DE genes in *M. bo*w*s*-infected macrophages in contrast to *M. tuberculosis* infection, mainly at 24 hpi (1,313 versus 904 genes respectively) and 48 hpi (2,271 versus 1,037 genes respectively) (Fig.3A, B). Comparison of the relative change with respect to control between *M. bovis-* and *M. tuberculosis*-Infected macrophages at each time-point revealed a statistically significant divergence in their responses at 48 hpi only associated with DE signatures from 703 genes (Fig.4A, Supp_VIII.csv). Analysis of the expression pattern of 576 of these genes with functional annotation across time revealed a greater magnitude of change in *M. bovis*-infected macrophages, where DE genes are up- or down-regulated to a higher extent in *M. bovis*-infected macrophages or not significantly changing at all in *M. tuberculosis*-infected macrophages (Fig.5C). Pathway enrichment analysis revealed association of *ABC transporters* with the divergent annotated gene set (Fig.5B); these are involved in cholesterol efflux from the cell, and manipulation of host cell cholesterol transport and metabolism has been documented in *M. tuberculosis* containing macrophages (82). A general dampening in the expression of cholesterol-associated genes was noted in *M. bovis*-infected macrophages at 48 hpi (Fig.S6B). Pathway enrichment analysis also associated *Cytosolic DNA-sensing* and *RIG-I-like receptor signalling* with the 576 divergent genes (Fig.5C). Type-I interferons have been associated with pathogenesis during *M. tuberculosis* infection and their production has been found dependent on the mycobacterial ESX-1 secretion system and the cytosolic sensing of extracellular *M. tuberculosis* DNA and subsequent cGAS-STING-dependent signalling (83-86).

**Figure 5:**
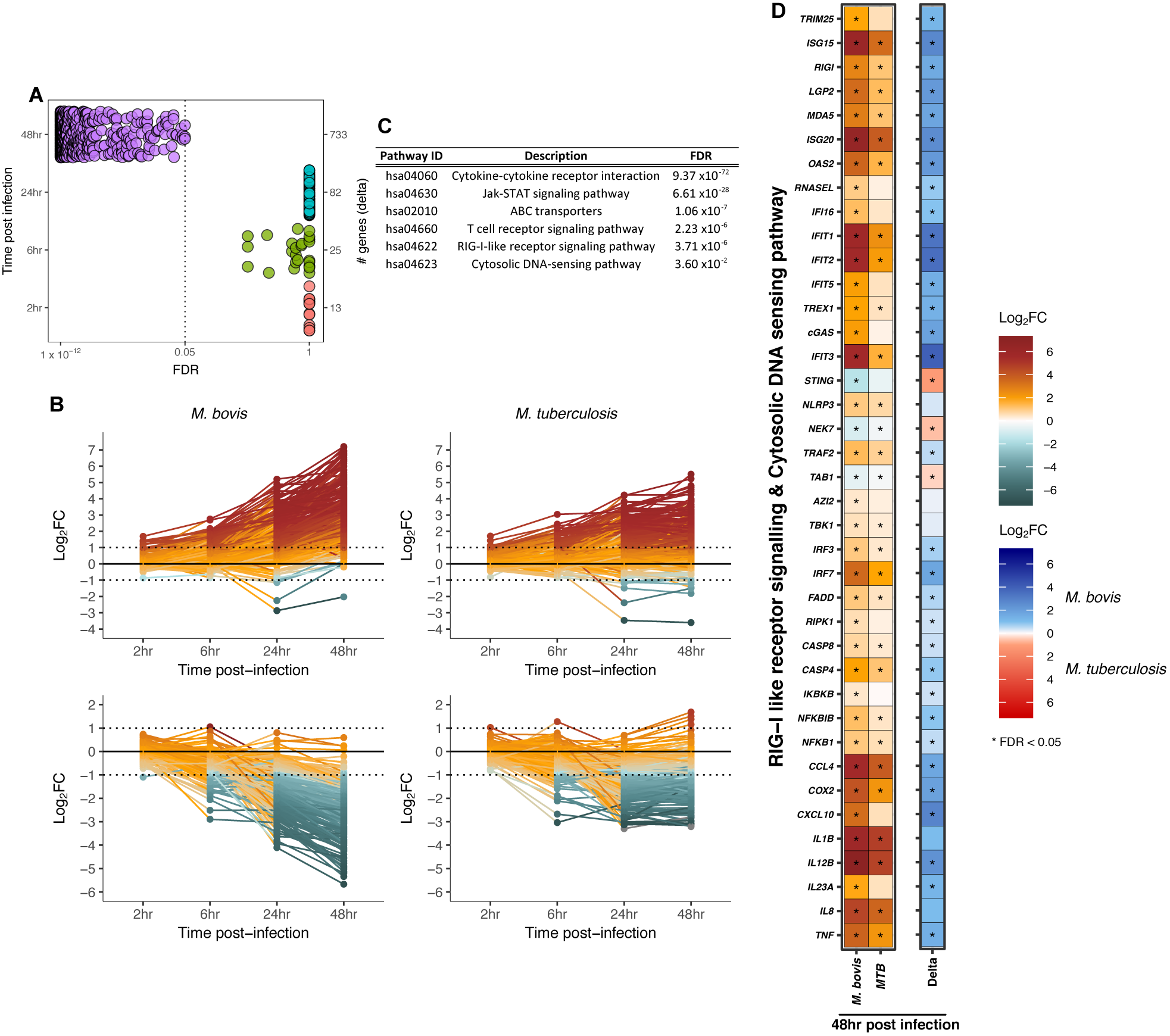
**A)** The number of genes (right y-axis) that are changing (|Log_2_FC| > 1) and that pass FDR threshold (FDR< 0.05) from the comparative analysis of *M. bovis-* or *M. tuberculosis-*infected macrophages in contrast to control macrophages and subsequently in contrast to the other infection series (delta comparison) at 2, 6, 24 and 48 hours post-infection (‘æ genes (delta)’) **B)** Line graphs represent those differentially expressed functionally annotated genes (*n =* 576) that exhibit a higher magnitude of change in *M. bovis*-infected macrophages versus *M. tuberculosis-Infected* macrophages in a positive manner (*n =* 323) (left and right top panel respectively) and in a negative manner [*n =* 253) (left and right bottom panel respectively) at 2, 4, 24 and 48 hours post-infection. **C)** Pathways that are enriched for 576 functionally annotated genes that exhibit divergent expression patterns in *M. bovis-* or *M. tuberculosis-Infected* macrophages at 48 hours post-infection (FDR < 0.05). **D)** The differentially expressed genes (|Log_2_FC| > 1, FDR < 0.05) associated with *RIG-l-like* and *DNA sensing signalling* pathways in bovine alveolar macrophages infected with *M. bovis* (blue) or *M. tuberculosis* (red) at 48 hours post-infection.

Overall, there is a stronger upregulation of genes encoding proteins involved in *RIG-I-like* and *DNA-sensing* signalling in *M. bovis*-infected macrophages in comparison to *M. tuberculosis-infected* macrophages at 48 hpi. These include genes encoding DNA sensors such as MB21D1/cGAS, MDA5, IFI16, and DDX58/RIG-I, antiviral and MAVS-TBK1 interacting protein IFIT3, serine/threonine kinase TBK1, and key interferon-I transcriptional regulators IRF3 and IRF7 that are all known to contribute to STING-dependent induction of type-I interferons (Fig.5D, Fig.6) (87-89). Furthermore, genes *LGP2, ISG15* and *TRIM25* that encode regulators of *DDX58/RIGI* gene expression are upregulated in *M. bovis*-infected macrophages at 48 hpi (90). Likewise, downstream of RIG-I, genes *IKBKB* and *1KB* are also expressed higher in *M. bovis*-infected macrophages along with *NFKB,* the gene encoding a key transcription factor that regulates the expression of inflammatory-related genes (89). Targets of NFKB such as *TNF, COX2, CXC40, MlPla, 118, 1112 and IL23a* are all expressed to a higher degree in *M. bovis-*infected macrophages with respect to *M. tuberculosis-infected* macrophages at 48 hpi (Fig.5D, Fig.6). A low level of reads mapped to the type I interferon genes *IFNAD* (orthologue of human *IFNA1*) and *IFNB1,* and only in a subset of animals at certain time points excluded these genes from DE analysis based on filtering criteria.

**Figure 6:**
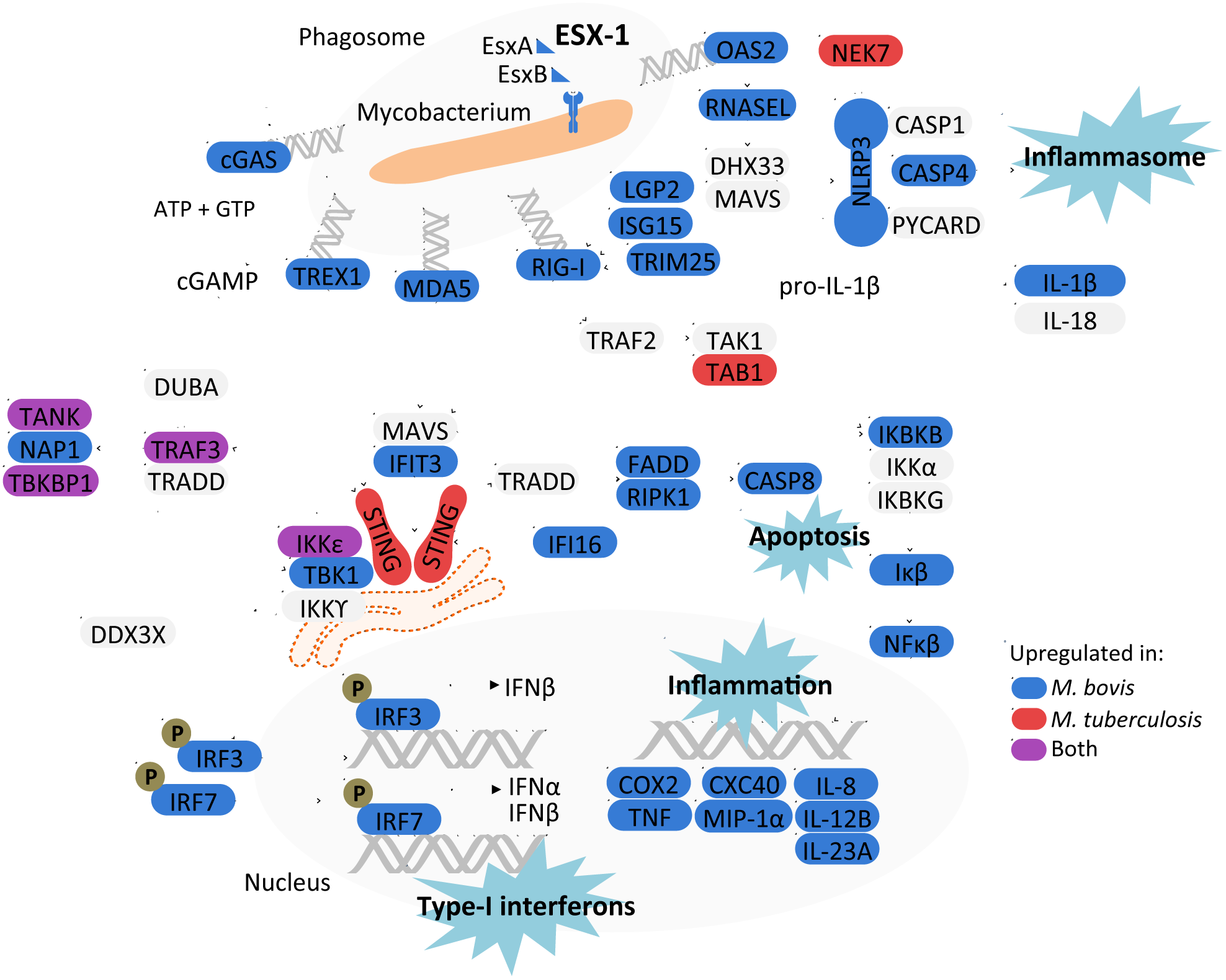
An overview of the DNA sensing and RIG-I signalling identified in this study 48 hours after infection of bovine alveolar macrophages with *M. bovis* and *M. tuberculosis.* Blue and red represents upregulation of the associated gene in either *M. bovis-* or *M. tuberculosis-’mfected* macrophages while purple represents upregulation of the associated gene in both infection models. The ESX-1 secretion system was found upregulated in *M. bovis* in comparison to *M. tuberculosis.*

Independent of RIG-I signalling, genes involved in *DNA sensing* such as *TREX1,* which encodes a 3’-5’ exonuclease that senses and degrades cytosolic DNA to prevent type I interferon production through the TBK1/STING/IRF3 pathway, and *OAS2* were also found expressed to a higher level in *M. bovis* containing macrophages (91). OAS2 is a double stranded RNA binding protein that generates 2’-5’- adenosine oligomers which activate RNase L resulting in the assembly of the NLRP3 inflammasome and IL-1b production (92, 93); genes *RNASEL, NLRP3* and non-canonical activation of the NLRP3 inflammasome *CASP4* were all found expressed to a higher level in *M. bovis*-infected macrophages 48hpi (94, 95) (Fig.4c, Fig.6). Taken together, these data highlight dissimilarity in the engagement between *M. bovis* and *M. tuberculosis* with the nucleic acid sensing system of the bovine macrophage, which in turn would influence downstream immune-related events, and ultimately infection outcome.

## Discussion

The data we present here provides significant insight on the molecular basis of host tropism between *M. tuberculosis* and *M. bovis* in the bovine host. Determining the differences between the transcriptional and translational profiles of these two hallmark mycobacterial strains highlighted variable expression of virulence-associated pathways while a divergent transcriptional response to infection with either species in bovine alveolar macrophages was observed 48 hpi. The ESX-1 secretion system of *M. tuberculosis* is linked to its phagosomal escape during infection, a process that is coupled to the triggering of DNA-sensing pathways in the cytosol of the host cell (83-86). In this study, we found that the ESX-1 secretion system was expressed to a higher level in *M. bovis* and a substantially stronger induction of DNA-sensing related pathways was seen in bovine alveolar macrophages infected with *M. bovis* versus *M. tuberculosis.* These data therefore suggest that *M. bovis* has a distinct engagement with the bovine immune system and might thus be better able to drive phagosome rupture and downstream immune signalling, leading to successful infection and ultimately disease.

In this study, we have taken advantage of both RNA-seq and SWATH mass spectrometry to compare both the global transcriptional and translational expression profiles of the human and bovine tubercle bacilli for definition of functional variation between the two species that may explain their exhibited host preference. Other studies have assessed in isolation either the transcriptome by microarray or the proteome by shotgun mass spectrometry (27, 28, 96, 97); the resolution afforded by both RNA-seq and SWATH mass spectrometry in comparison to previous studies has allowed for the most complete dataset for *M. bovis* to date and the most complete comparative dataset between the two pathogens. Although conducting a dual RNA-sequencing study may facilitate simultaneous assessment of the transcriptional response of both the host macrophage and invading mycobacteria during infection, this technique is limited with regards to the proportion of the bacterial-to-host transcriptome ratio in the resulting data and it does not allow for the accurate capture of both the global transcriptomic and proteomic dynamics of the mycobacteria during infection (98). As we aimed to investigate the early response of the macrophage to infection with both mycobacterial species, we believe the overall expression profiles measured in this study *in vitro* more realistically represent the bacterial phenotypes first encountered by the host cell. Lastly, we focused on *M. tuberculosis* H37Rv and *M. bovis* AF2122/97 as they are widely used reference strains and they have been previously used to demonstrate the attenuation of *M. tuberculosis* in the bovine host (1, 9).

The relatively small number of DE genes at both the RNA and protein level between *M. bovis* and *M. tuberculosis* highlights the close genetic relationship between the two pathogens. That being said, assessment of these DE genes supports our hypothesis that subtle genetic changes between the two species result in divergent phenotypes driven by differential expression of major virulence associated factors and pathways. We found that transcription factors PhoP, WhiB3 and DosR are significantly associated with the DE genes between the two species and these are functionally linked by processes that govern the adaptation of *M. tuberculosis* to the intracellular environment (53, 54). The PhoPR two-component system is important for *M. tuberculosis* infection and it has been suggested that mutations in PhoR attenuate animal-adapted *M. bovis* in humans (23, 99-105). PhoP regulates the production of virulence associated cell wall lipids and controls the expression of EspA, an ESX-1 secretion pathway related protein involved in the secretion of the major antigen EsxA/ESAT6 (55, 56, 70, 71). We found that the ESX-1 secretion system is expressed to a higher degree at both the RNA and protein level in *M. bovis* in comparison to *M. tuberculosis;* differences in ESX-1 secretion system expression between the two pathogens may be a consequence of a SNP in the promoter of the *whiB6* gene in *M. tuberculosis* H37Rv or attributed to attenuated PhoPR signalling in *M. bovis* (56). There is an emerging body of evidence showing that *M. tuberculosis* can rupture the phagosome membrane through the action of the ESX-1 secretion system and that the activation of cytosolic DNA-sensing pathways and the production of Type-I interferons is dependent on ESX-1 expression (83-86). Based on our data, we speculate that alternate transcriptional regulation between *M. tuberculosis* and *M. bovis* as a consequence of genetic variation may represent differential priming events in preparation for the initial interactions of both species with their respective host immune systems. For example, increased expression ESX-1 secretion system may facilitate faster escape of *M. bovis* from the phagosome into the cytosol in contrast to *M. tuberculosis,* triggering DNA-sensing pathways and increased type I interferon production (68, 85, 106).

To determine the impact of pathogen variation on host response, we conducted an experimental infection of primary bovine alveolar macrophages with *M. tuberculosis* and *M. bovis* and tracked the transcriptional response to infection. The bovine alveolar macrophage response to infection with either pathogen was strikingly similar over the first 24 hours of infection. Notably, a “core” macrophage response displayed enrichment for differentially expressed genes involved in pathogen recognition, innate cell signalling, and cytokine and chemokine production illustrating the initiation of host innate defence mechanisms in response to infection with *M. bovis* and *M. tuberculosis.* One of the most striking observations is that divergence in macrophage gene expression profiles between *M. bovis* and *M. tuberculosis* infections only occurs after 24 h, with *M. bovis* infection eliciting a stronger response in comparison to *M. tuberculosis.* At 48 hpi, enrichment for *DNA sensing* was found for 576 annotated genes that show divergent expression patterns between the two infection models. The innate immune system detects exogenous nucleic acid within the cell through pattern recognition receptors (PRRs) that include Absent in Melanoma 2 (AIM2)-like receptors (ALRs) with Pyrin and HIN domains (PYHIN proteins), *e.g.* IFI16 (107-109). Other DNA-sensing proteins include cytosolic RIG-I-like receptors (RLR), (*e.g.* RIG-I, MDA5, LGP2), exonucleases, synthetases, and cyclic GMP-AMP synthases (*e.g.* TREX1, OAS2 and cGAS) (83, 91, 93, 110, 111). A stronger transcriptional induction of genes associated with cGAS-STING dependent signalling was seen in macrophages infected with *M. bovis* including *MB21D1/cGAS* and downstream effectors *TBK1* and *IRF3* (Fig.6). cGAS has a central role during *M. tuberculosis* infection; 48-72 hpi cGAS senses *M. tuberculosis* in the host cell cytosol and in turn signals through STING to drive type-I interferon production (Fig.6) (68, 83-86, 112). Surprisingly, the cGAS-STING axis was not the only PRR pathway found upregulated during mycobacterial infection as *RIG-I like signalling* pathway was also observed to be enriched at 48 hpi, with genes encoding TREX1, OAS2, and RLR receptors RIG-I, MDA5 and LGP2 also found expressed to a higher level in *M.* bovis-infected macrophages. As we hypothesised that the variation in host response at 48 hpi, and ultimately host tropism, is driven by differential expression of virulence factors between *M. bovis* and *M. tuberculosis,* the identification of an increase in expression of DNA-sensing related pathways in *M. bovis*-infected macrophages at 48 hpi coincides with the differential expression of the ESX-1 secretion system between the two pathogens. A further role for the ESX-1 secretion system in host-pathogen interactions is described through the activation of the host NLRP3 inflammasome and the production of IL-lb (113-116). Transcriptional signals associated with the NLRP3 inflammasome were higher in *M. bovis*-infected macrophages at 48 hpi along with the increased expression of *CASP4,* an NLRP3 inflammasome activator, which has a central role in mediating the response to *Legionella, Yersinia* and *Salmonella* bacterial infection in primary human macrophages and that has been found upregulated in the necrotic granuloma model of mice and lymph nodes of TB patients (92, 117-119). Altogether, these data therefore suggest that not only does mycobacterial infection in the bovine macrophage trigger an increase in the transcription of the cGAS/STING/IRF3 pathway previously characterised as responsible for type-I interferon production during *M. tuberculosis* infection, it also triggers alteration in the transcription of genes encoding auxiliary DNA sensing RLR receptors including RIG-I, MDA5 and TREX1, that likewise converge to signal through the STING complex (Fig.6). Furthermore, the data highlights that *M. bovis* drives a stronger transcriptional response in the aforementioned pathways in the bovine alveolar macrophage at 48 hpi in comparison to *M. tuberculosis,* again highlighting a distinct relationship between the bovine pathogen and the bovine host.

Altogether, the upregulation of the ESX1 secretion system at both the RNA and protein level in *M. bovis* with the observed upregulation of DNA-sensing pathways and the NLRP3/IL-1b pathway in *M. bovis*-infected macrophages suggests that the expression level of virulence factors, rather than the presence or absence of them between the highly related *M. bovis* and *M. tuberculosis,* drives divergent host responses and influences infection outcome overall. Indeed, the idea of a ‘perfect balance’ with regards to the expression of mycobacterial virulence factors is reflected in the findings that production of IFN-β1 in monocyte-derived macrophages is strain dependent amongst the *M. tuberculosis* lineages (120). That being said, we cannot disregard that genetic differences between the bovine and human host may play a factor. As the innate immune response in different mammals can vary, diversity in the expression and structure in key innate immune genes and engagement with pathogen factors must play major roles in host specificity and the outcome of pathogen encounter (121). In this regard, it is interesting to note that the bovine PYHIN locus contains only *IFI16* (bovine PYHIN) and cattle are the only mammals to date found to encode a single member of PYHIN protein family; in contrast, humans have four genes, and mice 13 genes (122). Furthermore, polymorphisms in *NLRP3* have been found to influence host susceptibility to *M. tuberculosis* infection, its induction is associated with the mycobacterial ESX-1 secretion system, and bovine and human NLRP3 proteins share 83% sequence identity (123). Further comparative studies of human and bovine genetics will ultimately aid in unravelling the complex differential host response to infection with both pathogens. Moreover, we cannot overlook other virulence-associated factors besides the ESX-1 secretion system that differ in expression or sequence between *M. bovis* and *M. tuberculosis.* Indeed, antigens MBP83 and MBP70, that show constitutive upregulation in *M. bovis* versus *M. tuberculosis* and pathways such as ESX-3 and Mce-1 were also found differentially expressed between the two pathogens at both the transcriptional and translational level during *in vitro* growth in this study (Supp_III.xlsx)(47, 124).

In conclusion, we found that *M. tuberculosis* H37Rv and *M. bovis* AF2122/97 induce divergent responses in infected bovine alveolar macrophages, a consequence of the differential expression of key mycobacterial virulence-associated pathways. Our work demonstrates the specificity of mycobacterial host-pathogen interaction and indicates how the subtle interplay between the phenotype of the invading mycobacteria and the subsequent host response may underpin host specificity amongst members of the MTBC.

## Acknowledgements

We would like to thank Dr. Christina Ludwig for help with the SWATH mass spectrometry measurements and Dr. Maximiliano Gutierrez for valuable input and discussion. We gratefully acknowledge funding from Science Foundation Ireland through SFI Investigator Awards 08/IN.1/B2038 and 15/IA/3154; the European Commission’s H2020 program grant number 643381 (TBVAC2020); Wellcome Trust PhD awards 097429/Z/11/Z (K.R-A.) and 102395/Z/13/Z (A.S.). The work was further supported by funding from SystemsX.ch/TbX (R.A.) and a research grant from Institut Mérieux (R.A.).

---

**Figure S1:**
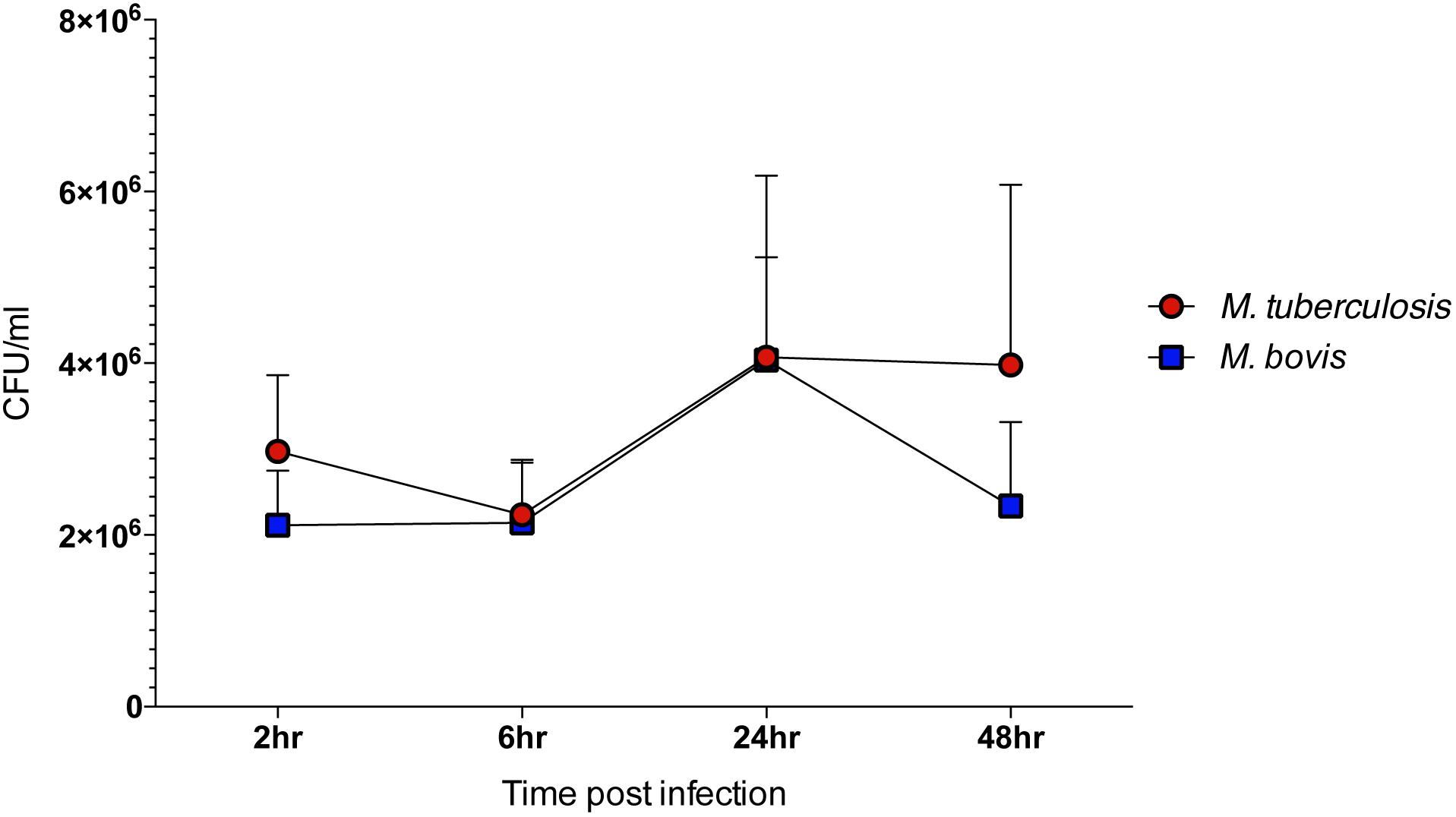
The number of colony forming units (‘CFU/ml’) recovered from bovine alveolar macrophages infected with *M. bovis* (blue) or *M. tuberculosis* (red) at 2, 6, 24 and 48 hours post-infection. (Error bars represent standard error of the mean, *n =* 6)

**Figure S2:**
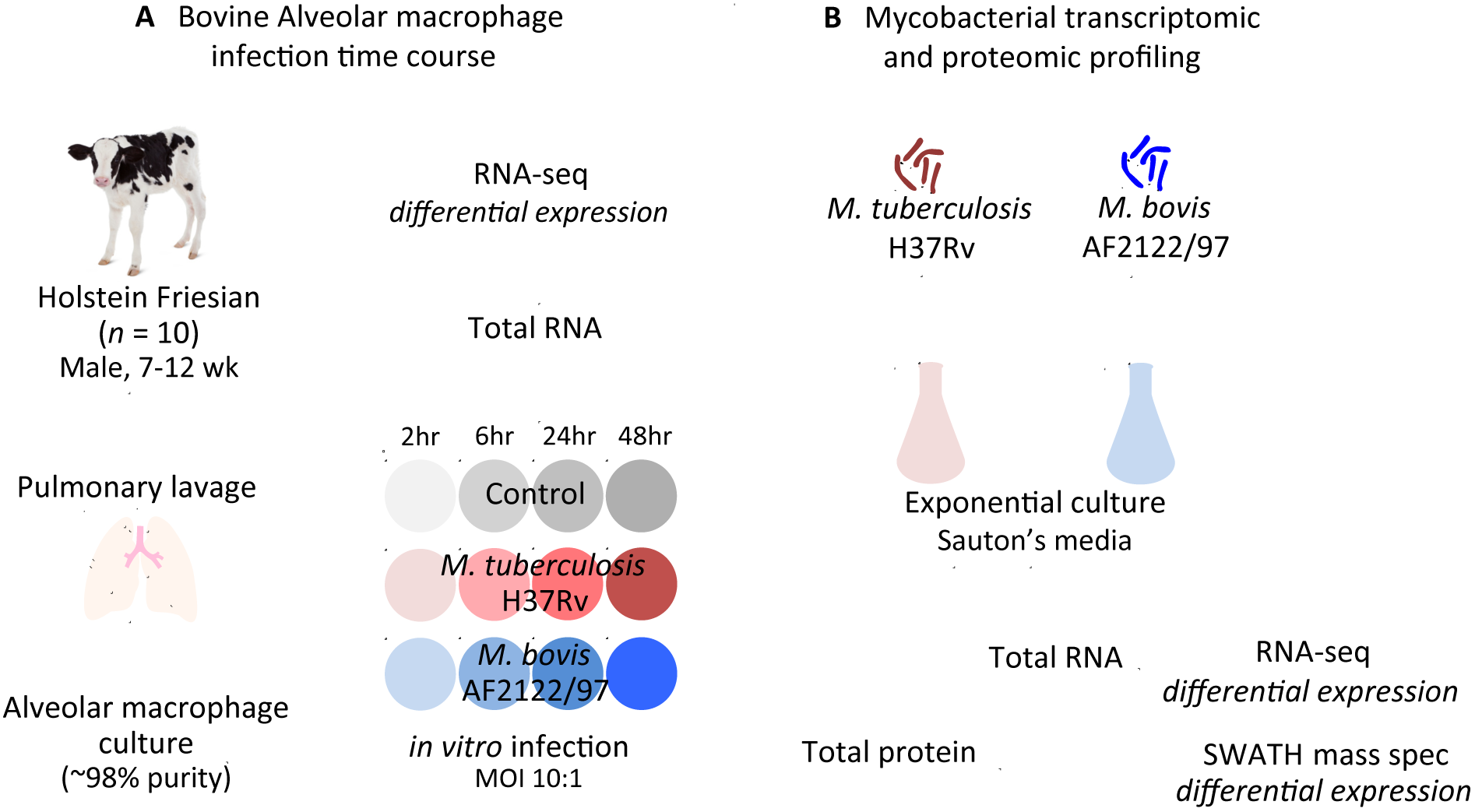
Overview of experimental design for the **A)** bovine alveolar macrophage infection time course and **B)** mycobacterial transcriptomic and proteomic profiling performed for this study.

**Figure S3:**
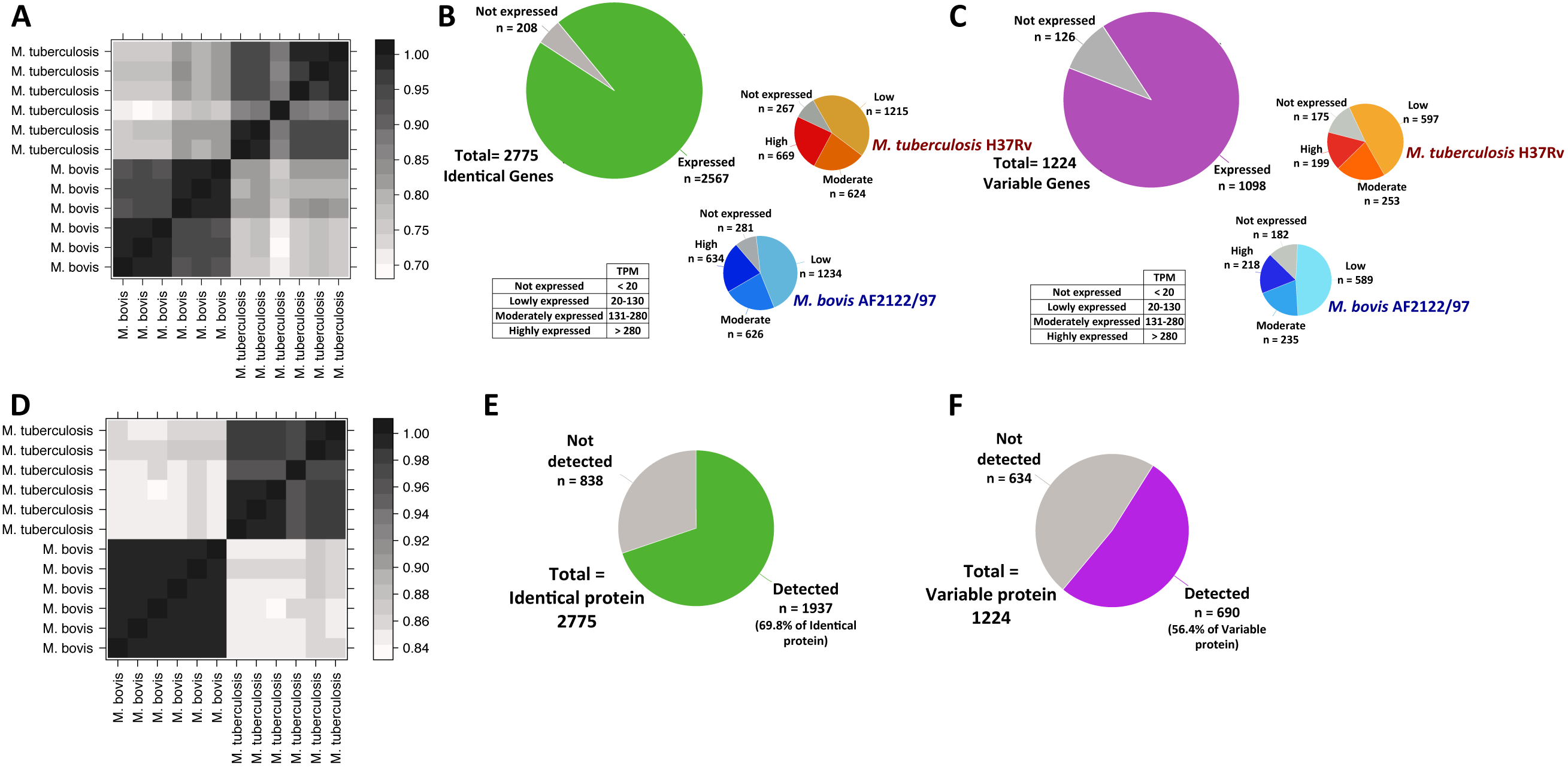
**A)** Pearson correlation plot of reads mapped to 2,775 Identical genes (100% conserved in length and amino acid sequence between the two species) in the six *M. bovis* and six *M. tuberculosis* RNA-seq datasets. Pie charts representing the proportion of B) Identical (100% conserved in length and amino acid sequence between the two species, green) and C) Variable genes (< 100% conserved in length and amino acid sequence between the two species, purple) detected and not detected across *M. bovis* and *M. tuberculosis* RNA-seq datasets. RNA expression values (Transcripts per Million (TPM)) were calculated for each gene and gene expression within either species was categorised into not expressed (<20 TPM), lowly expressed (20-130 TPM), moderately expressed (131-280 TPM) and highly expressed (>280 TPM). **D)** Pearson correlation plot of the intensity values of the 2,627 identified proteins in the six *M. bovis* and six *M. tuberculosis* SWATH MS datasets. Pie charts representing the expression of **E)** 2775 Identical genes (100% conserved in length and amino acid sequence) and **F) 1**224 Variable genes (< 100% conserved in length and amino acid) in *M. bovis* and *M. tuberculosis* detected by SWATH MS.

**Figure S4:**
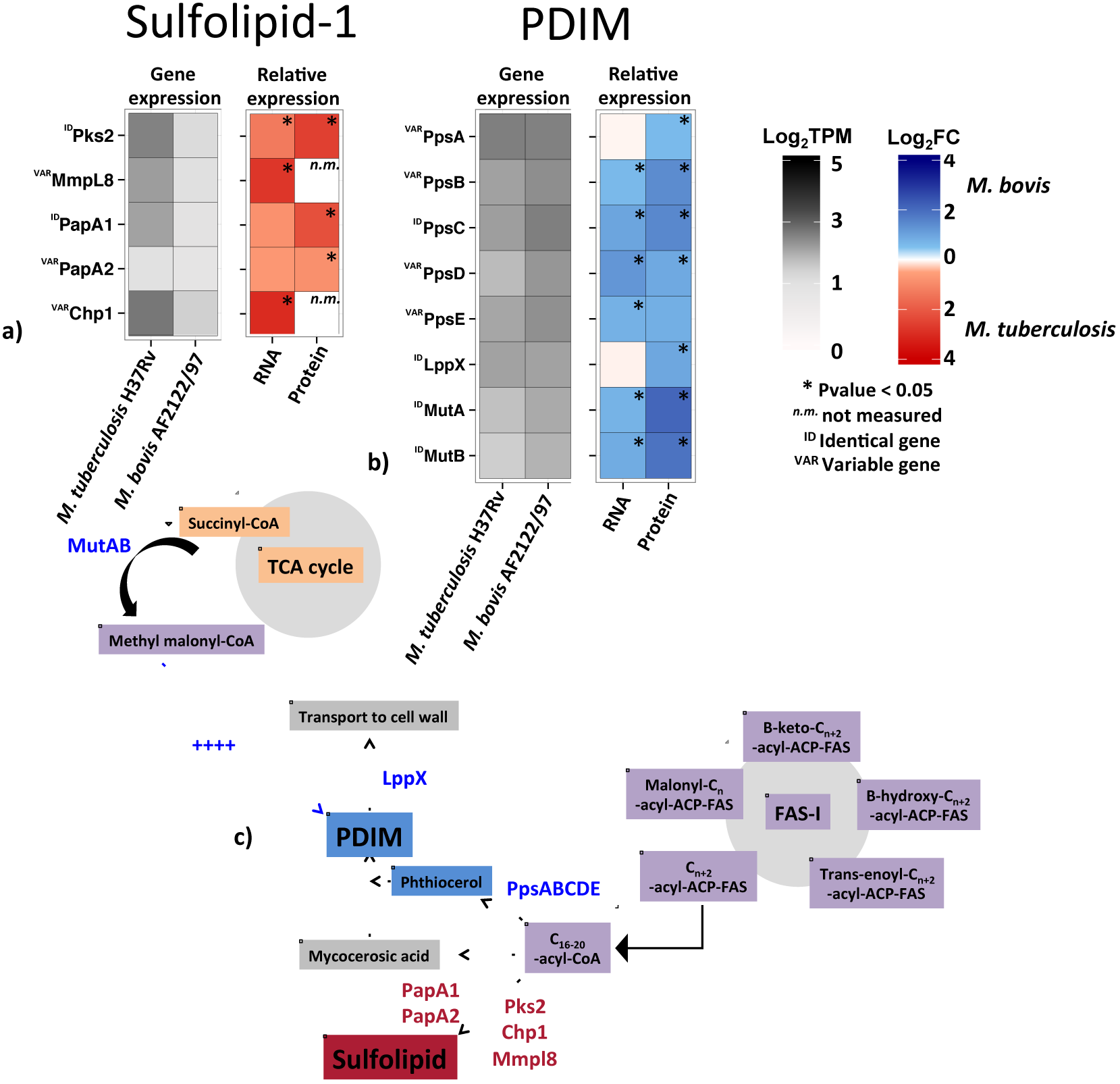
The expression of the **a)** sulfolipid-1 (SL-1) and **b)** phthiocerol dimycocerosate (PDIM) synthesis associated genes at the RNA and protein level in *M. tuberculosis* (red) and *M. bovis* (blue). The expression of each gene (“Gene expression”) is presented as LogioTPM at the RNA level while the relative expression (“Relative expression”) between the two species is presented as log2FC. Those genes that change significantly at the RNA and protein level (FDR < 0.05) are denoted (‘*’). **c)** Diagrammatic overview of the SL-1 and PDIM biosynthesis pathways in *M. tuberculosis.* Blue represents the genes in b) that are upregulated in *M. bovis* in contrast to *M. tuberculosis* and red represents the genes in a) that are upregulated in *M. tuberculosis.*

**Figure S5:**
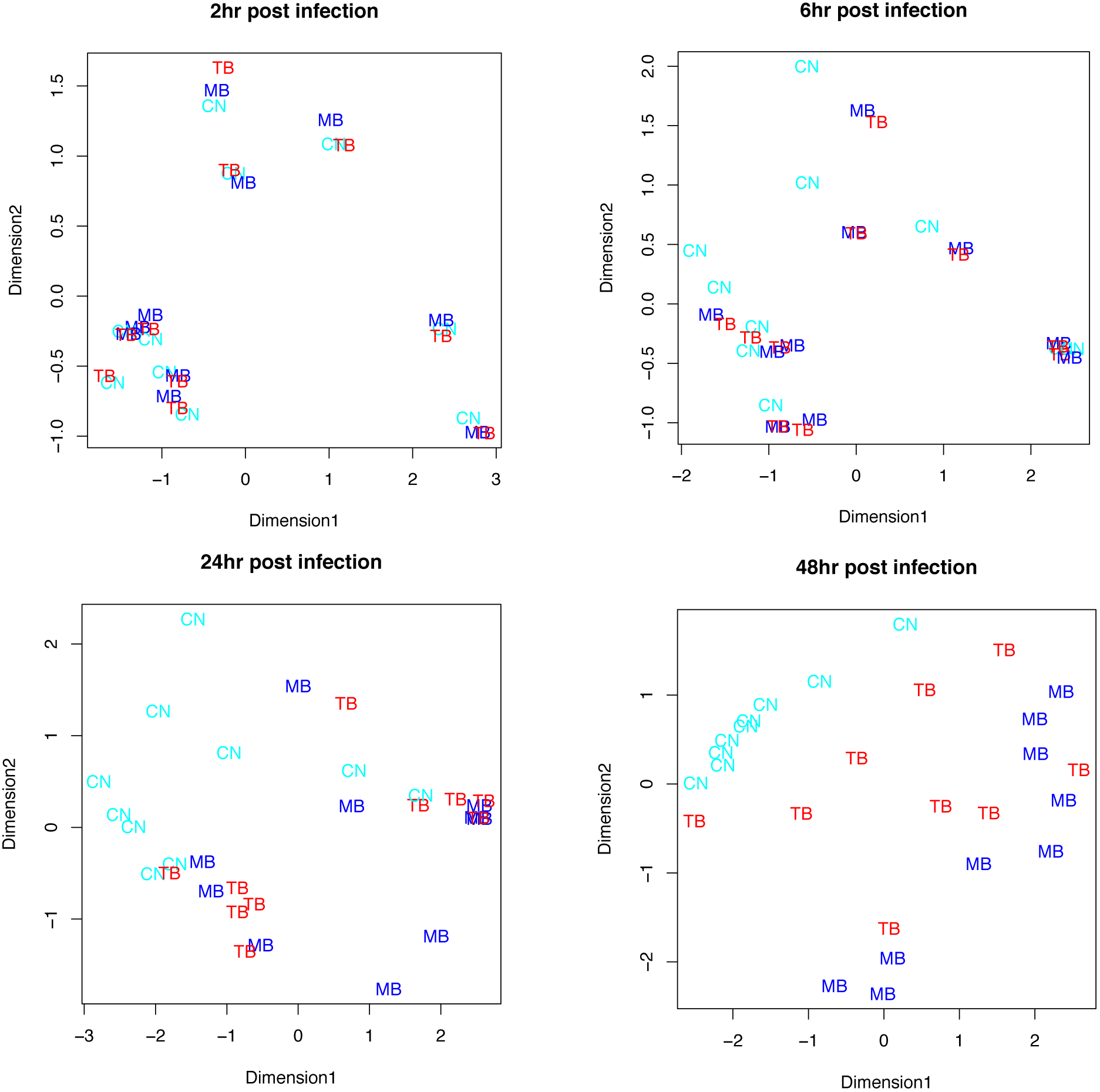
Multidimensional scaling plots of the RNA-seq expression data for individual of bovine alveolar macrophages infected with *M. bovis* (‘MB’, blue), *M. tuberculosis* (‘TB’, red) or none (‘CN’, cyan) at 2, 6, 24 ad 48 hours post-infection.

**Figure S6:**
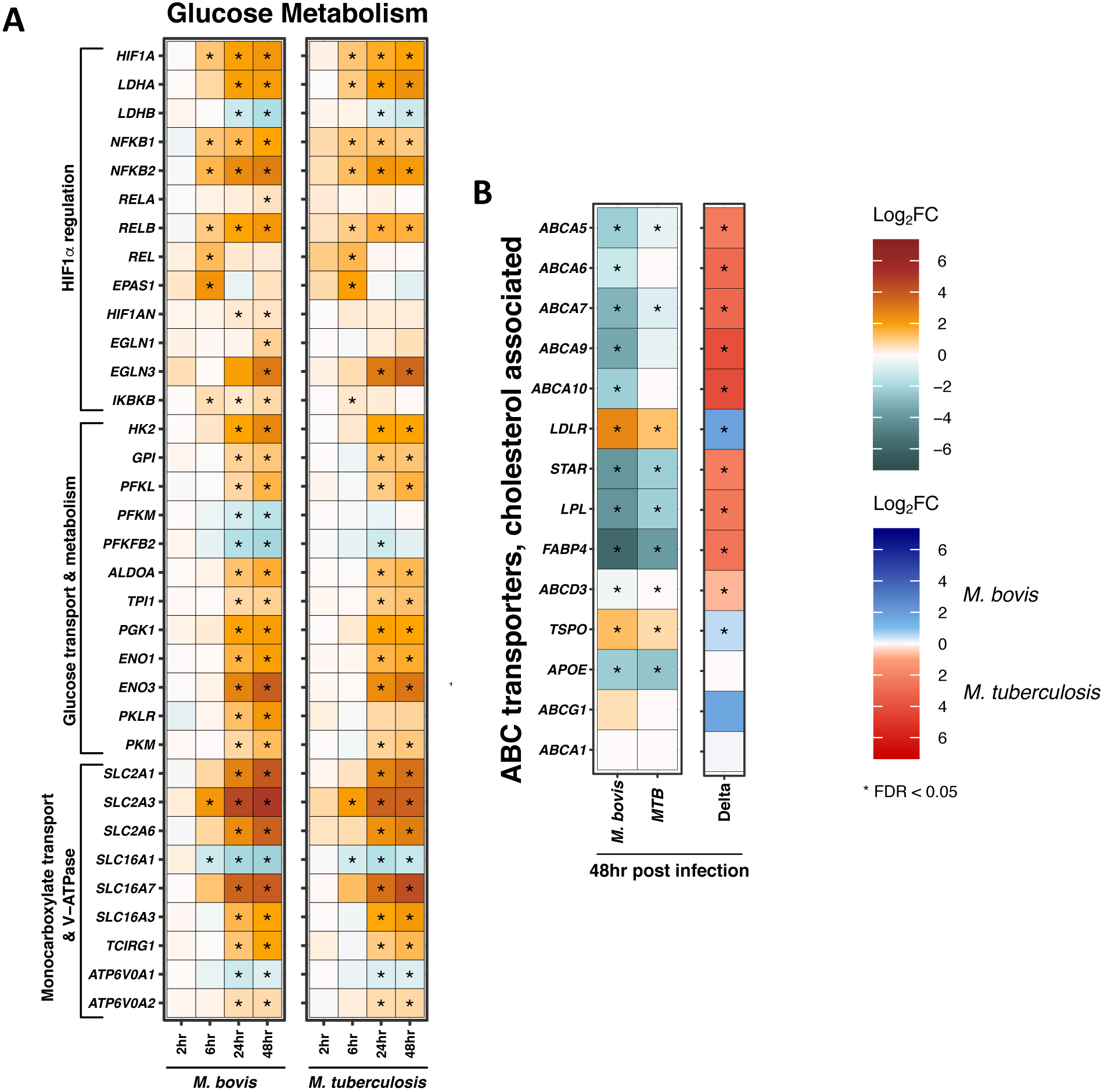
The expression of differentially expressed genes (|Log_2_FC| > 1, FDR < 0.05 (‘*’) associated with **A)** glucose metabolism in bovine alveolar macrophages infected with *M. bovis* or *M. tuberculosis* (“MTB”) at 2, 6, 24 and 48 hours post-infection and **B)** cholesterol-associated transport in bovine alveolar macrophages at 48 hours post-infection. Delta comparison shows genes upregulated in *M. bovis* in blue and *M. tuberculosis* in red.

